# High density of white-faced capuchins (*Cebus capucinus*) and habitat quality in the Taboga Forest of Costa Rica

**DOI:** 10.1101/692293

**Authors:** Elizabeth Tinsley Johnson, Marcela E. Benítez, Alexander Fuentes, Celia R. McLean, Ariek B. Norford, Juan Carlos Ordoñez, Jacinta C. Beehner, Thore J. Bergman

## Abstract

Across the globe, primate species and habitats are threatened by human activity. This is especially true for species found in tropical dry forests, which are widely distributed and comprise diverse habitats that remain largely unprotected. Evidence suggests that some primate species endemic to tropical dry forests may be more sensitive to anthropogenic disturbance than others, but our ability to predict primate abundance in the face of disturbance also depends on the specific variables for each site. Here, we consider the factors that explain the high density of white-faced capuchins (*Cebus capucinus*) found in the Taboga Forest, Costa Rica, a relatively small fragment of tropical dry forest surrounded by agricultural fields. Our analyses suggest that, for capuchins (and potentially for mantled howler monkeys, *Alouatta palliata*), the size and disturbance of a forest fragment may matter less than the composition and availability of key resources, like above-ground water. Group sightings for both species were higher near permanent water sources, but group sightings did not vary between edge and interior forest. These findings help explain why some primate species can flourish even alongside anthropogenic disturbance and thus carry important implications for conservation efforts. Smaller forest fragments, like Taboga, may be able to support high densities of some species because they provide a mosaic of habitats and key resources that buffer adverse ecological conditions. Future studies will assess the extent to which primates in the Taboga Forest rely on the canals versus the river and will consider how the high density of capuchins in Taboga influences ranging patterns, home range overlap, and the frequency and intensity of intergroup encounters.

**RESEARCH HIGHLIGHTS:**

- Here we introduce a new white-faced capuchin study site in the Taboga Forest, Costa Rica, a fragmented tropical dry forest.
- Forest fragments like Taboga may support high primate densities because they provide a mosaic of habitats and key resources.

## INTRODUCTION

The majority of primate species across the globe are either under threat of extinction or experiencing population declines (Estrada et al., 2018). Non-human primate densities tend to decrease in unprotected areas, yet non-human primates (hereafter, “primates”) can nevertheless still flourish in areas of human activity, suggesting both a vulnerability and a resilience to anthropogenic disturbance (Cavada, Barelli, Ciolli, & Rovero, 2016). Some human activities (e.g., those contributing to climate change) tend to be more disruptive than others (e.g., habitat fragmentation), but different primate species also exhibit differential responses to each threat. Some species show remarkable behavioral flexibility and quickly adjust to new habitats, while others get pushed closer to extinction (Kulp & Heymann, 2015; Laurance et al., 2007; Ries, Fletcher, Battin, & Sisk, 2004). This variation is likely due to a number of factors, from species-specific characteristics (i.e., dietary breadth) to habitat characteristics (i.e., species richness). For example, in Bornean forests tree density predicts primate species richness much better than the degree of habitat disturbance does (Bernard et al., 2016). Understanding how species and habitat characteristics together contribute to resilience is critical for effective conservation efforts.

One key primate habitat that remains relatively understudied is the tropical dry forest. Tropical dry forests are widely distributed and diverse habitats that simultaneously support a number of endemic species while also experiencing significant anthropogenic disturbance (Dryflor et al., 2016; Miles et al., 2006). Despite warnings about the vulnerability of these habitats (e.g., Janzen, 1988), tropical dry forests worldwide remain unprotected and understudied (Dexter et al., 2018). For example, over 90% of the tropical dry forests in North and Central America are vulnerable to anthropogenic disturbance (Miles et al., 2006). However, a variety of primate species are found in tropical dry forests, with some even continuing to flourish in fragments. Most notably, white-faced capuchins (*Cebus capucinus*), howler monkeys (*Alouatta palliata*), and spider monkeys (*Ateles geoffroyi*) are common sympatric species, yet they demonstrate markedly divergent responses to fragmentation and other forms of anthropogenic disturbance (e.g., Williams-Guillén, Hagell, Otterstrom, Spehar, & Gómez, 2013).

Understanding how different species respond to anthropogenic disturbance has important implications for conservation and reforestation efforts. This is especially true when it comes to tropical dry forests, which were once the predominant forest type on the west coast of Central America (Gillespie, Grijalva, & Farris, 2000). While spider monkeys are rarely found in small fragments (instead requiring large territories to support their frugivorous diet: Williams-Guillén et al., 2013), both capuchins and howlers can thrive there. As large-bodied omnivores, capuchins, in particular, can opportunistically exploit a broad array of plants and animals (Ford & Davis, 1992; Rose, 1994; Panger et al., 2002; Perry 2012) and can adapt to anthropogenic disturbances that threaten many other species (i.e., showing neutral or even positive edge effects: Bolt et al., 2018 [*C. capucinus*]; surviving in fragmented habitats: Lins & Ferreira, 2019 [*Sapajus flavius*]; Cunha, Vieira, & Grelle, 2006 [*C*. sp.]). However, certain factors may constrain capuchin densities. For example, low capuchin densities in Nicaraguan forest fragments have been attributed to human hunting, poaching, and the pet trade (Williams-Guillén et al., 2013). In addition, capuchin populations in dry forests, where rainfall is scarce for months at a time, may be constrained by access to reliable above-ground water (Fedigan & Jack, 2001). Howler monkeys, in contrast, may be less reliant on above-ground water sources because of the high water content in the leaves in their diet (Fedigan & Jack, 2001). Moreover, howlers are expected to do well in disturbed habitats in general, and data tend to support this (i.e., showing neutral or positive edge effects: Bolt et al., 2018 [*A. palliata*]; Lenz, Jack, & Spironello, 2014 [*A. macconelli]*; surviving in fragmented habitats: Asensio, Arroyo-Rodríguez, Dunn, & Cristóbal-Azkarate, 2009 [*A. palliata mexicana*]; Boyle & Smith, 2010 [*A. macconelli*]). Despite this resilience under certain circumstances howler monkeys can still be negatively impacted by fragmentation (Arroyo-Rodríguez & Dias, 2010; Horwich, 1998). For example, a 1976 census of howlers at Taboga suggested that the population was in decline, if not at a nadir (Heltne, Turner, & Scott, 1976).

Here, we studied plant and wildlife abundance in the Taboga Forest (hereafter, “Taboga”) of Costa Rica. Taboga presents an ideal opportunity to understand primate abundance in relation to habitat quality for a number of reasons. First, the Taboga Forest has an extremely high density of capuchins compared to other forests in the region (see Table 2). Second, the forest is relatively small (516 hectares) and irregularly shaped with a high proportion of edge to interior (almost 40%: Fig. 1). Third, the forest is dissected by a series of canals used in irrigation and has been completely surrounded by sugar cane and rice farmland for decades. Therefore, we are able to look at the long-term impacts of two types of human disturbance: habitat fragmentation and year-round, artificial water sources. This is particularly important because Taboga is a tropical dry forest, where the dry season would normally limit the viability of many fauna that depend on above-ground water.

**Table 1.** Group size and composition for three habituated white-faced capuchin groups

| Group | Adult males | Adult females | Subadults and juveniles | Infants | Total |
| --- | --- | --- | --- | --- | --- |
| Mesas | 2 | 4 | 6 | 4 | 16 |
| Tenori | 3 | 4 | 7 | 3 | 17 |
| Palmas | 5 | 8 | 12 | 4 | 29 |

**Table 2.** Group size and density comparison for this (“Taboga”) and other white-faced capuchin sites.

| Site | Description | Individuals | km2 | Individuals / km2 | Source |
| --- | --- | --- | --- | --- | --- |
| Lomas | Mean (3 groups) | 29.00 | 3.64 | 7.97 | Vogel, 2004 |
| Lomas | Total population | 216.00 | 36.99 | 5.84 | Perry, pers comm |
| Palo Verde | Total population | - | - | 9.40 | Panger et al., 2002 |
| Santa Rosa | Mean (7 groups) | 20.10 | 1.98 | 10.15 | Campos et al. 2014, Fedigan & Jack 2012 |
| Santa Rosa | Total population | 673.00 | 57.70 | 11.66 | Campos, pers comm |
| BCI | Mean (4 groups) | 15.00 | 1.16 | 12.93 | Crofoot, 2007 |
| BCI | Total population | 300.00 | 15.00 | 20.00 | Crofoot, 2007 |
| BCI (after 2010 crash) | Total population | 84.00 | 15.00 | 5.60 | Milton & Giacalone, 2014 |
| Taboga | Mean (3 groups) | 21.00 | 0.97 | 21.65 | Tinsley Johnson et al., this publication |
| Taboga | Total population | 187.00 | 5.17 | 36.17 | Tinsley Johnson et al., this publication |

**Figure 1.**
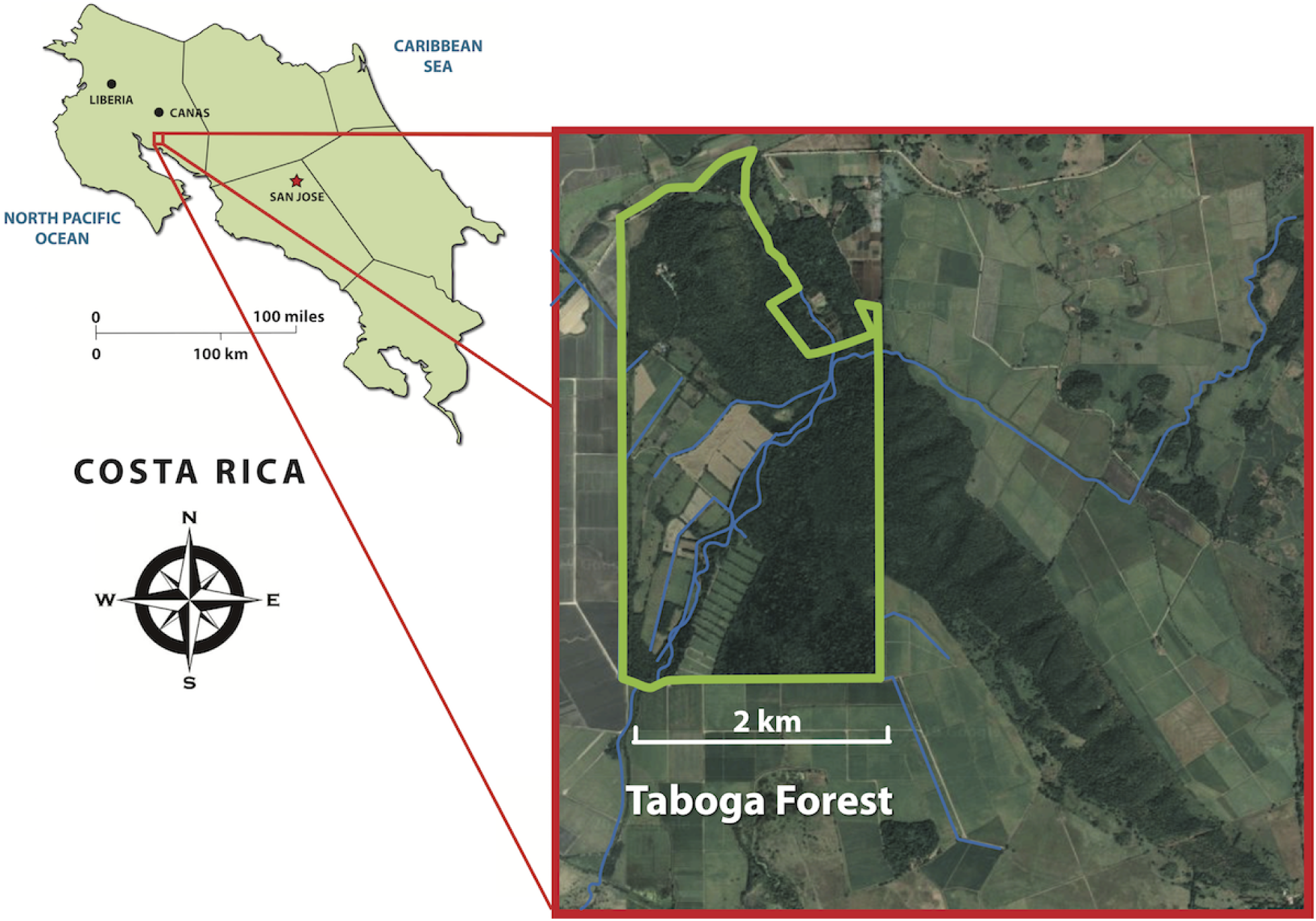
Location of the Taboga Forest in Costa Rica. The official Taboga Forest Reserve boundary is within this larger reserve held by the Universidad Técnica Nacional (see Fig. 3a). However, for simplicity, we refer to this entire area as the Taboga Forest or “Taboga”.

One of our primary goals is to estimate the density of white-faced capuchins in the forest (as well as the mantled howlers to serve as a comparison species). Specifically, we address the following questions: (1) What is the density of capuchins in Taboga, and how does this compare to other long-term sites with white-faced capuchins? To answer this, we first compare group size and home range size of known groups across long-term capuchin sites. (2) Does the large amount of edge habitat contribute to capuchin abundance at Taboga? To answer this, we next compare the plant composition of edge and interior forest. We predict higher species richness, mean diameter at breast height (DBH), and canopy coverage in interior compared to edge forest (Bolt et al., 2018). Third, we expect to find significant seasonal effects on canopy coverage, with lower coverage during the dry season. However, because capuchins have been shown to have neutral or even positive edge effects (e.g., Bolt et al., 2018), we expect to find no significant differences in species richness and mean DBH for tree species associated with capuchins (i.e., that capuchins use for food or fur-rubbing). Fourth, because of the long-term anthropogenic disturbance around Taboga, we expect to find significant differences in the species richness and DBH for indicator tree species associated with the first stage of forest succession in tropical dry forests (Kalacska et al., 2004).

Finally, we ask: (3) Do forest characteristics differentially affect capuchin as compared to howler abundance? We expect that both howler and capuchin monkeys will show neutral edge effects (Bolt et al., 2018). However, because Taboga is a tropical dry forest, we expect to find higher primate abundance within 100 m of reliable water sources (i.e., rivers and large canals), but this should mainly apply to capuchin groups, as howler monkeys are less dependent on above-ground water (Fedigan & Jack, 2001).

## METHODS

### Study site and subjects

We conducted this study at the Capuchins at Taboga research site, established in June 2017 in la Reserva Forestal Taboga (the Taboga Forest Reserve; i.e., Taboga) in Guanacaste province, Costa Rica. The reserve was established in 1978 and contains 296 hectares of largely tropical dry forest (∼3 × 4 km), representing an important piece of the fragmented biological corridor connecting the Guanacaste Mountains to the Tempisque River Basin (Fig. 1). The Universidad Técnica Nacional (UTN) of Costa Rica operates an experimental farm of 702 hectares that encompasses the reserve along with agricultural land. The UTN farm consists of irrigated land dedicated to the cultivation of sugarcane (100 hectares), rice (30 hectares), and grass for cattle (4.5 hectares). There is also a tilapia farm and research center as well as a water research laboratory. As such, Taboga is characterized by distinct forest edges (i.e., farm land and roads) as well as more transitional or “natural” forest edges (i.e., canals and rivers). The reserve is almost exclusively bordered by sugarcane and rice fields, aside from a 2 km perimeter that borders private forested land and 1 km bordering public forested land.

Taboga is largely characterized by seasonally dry tropical forest, a highly threatened ecosystem that features a closed canopy and seasonal deciduousness (Janzen, 1988; Miles et al., 2006). In addition to the dry forest, there are also riparian, semi-deciduous forests around the river and a palm forest dominated by the native species *Attalea rostrata*, part of which becomes inundated during the wet season. The area experiences two distinct seasons (Fig. 2): a hot, dry season from Nov-Apr (mean daily maximum temperature = 35.38 °C +/− 0.20 SE; mean daily rainfall = 0.66 mm +/− 0.27 SE) and a cooler wet season from May-Oct (mean daily maximum temperature = 32.57 °C +/− 0.21 SE; mean daily rainfall = 8.93 mm +/− 1.09 SE). Mean daily minimum temperatures remain consistent throughout the year (dry season: 26.25 °C +/− 0.10 SE; wet season: 25.46 °C +/− 0.11 SE). The river provides fresh water year-round and many of the canals used by the UTN farm for irrigation are consistently full during the dry season (Fig. 3C).

**Figure 2.**
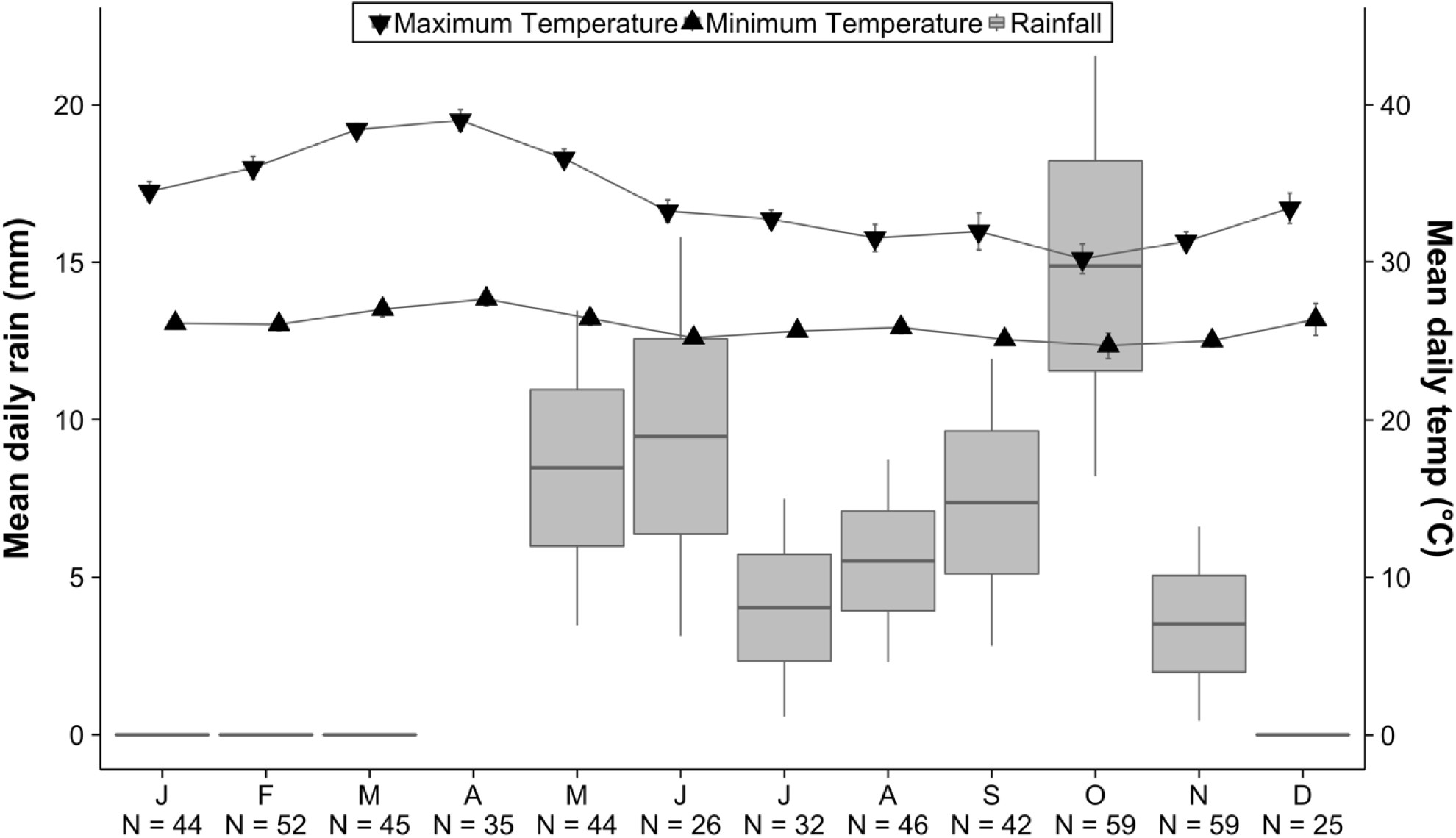
Temperature (black triangles) and rainfall (grey boxes) data from the Taboga Forest from Jul 2017 - May 2019. Numbers along the x-axis indicate the number of days of weather data measured per month.

**Figure 3a-d.**
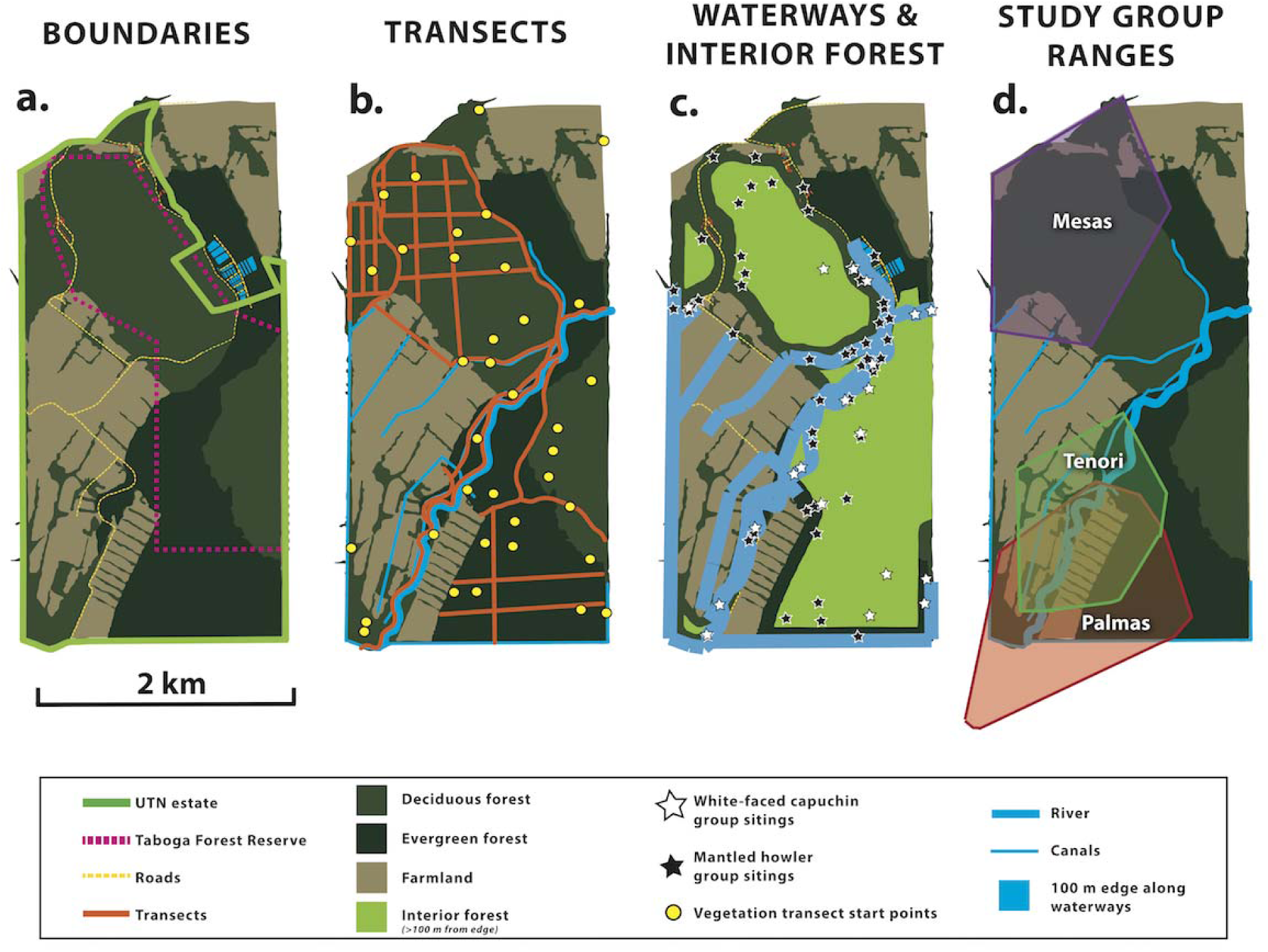
Maps of **(a)** the UTN Estate and the Taboga Forest Reserve (together, the “Taboga Forest”), with overlays displaying: **(b)** both primate and vegetation transect locations, **(c)** 0.1 km buffer zones for forest edges and year-round water sources, and **(d)** home ranges for three groups of wild white-faced capuchins.

The Capuchins at Taboga research project (directed by Thore Bergman, Jacinta Beehner, Marcela Benítez, Elizabeth Tinsley Johnson) is focused on the behavioral biology, endocrinology, and cognition of wild white-faced capuchins (*C. capucinus*). Note that the taxonomy of Central American capuchins is in flux and some authors refer to Costa Rican capuchins as *C. capucinus imitator* (Ruiz-Garcia et al., 2012; Lynch Alfaro et al., 2014; Melin et al., 2017; Hogan et al., 2018). However, we prefer the more general name *C. capucinus* because the defining feature of the capuchins first described as “imitator” is a frontal tuft in females, something that is clearly absent in our populations (Thomas, 1903). Our research group has identified at least 12 distinct capuchin groups in the reserve since July 2017. We collect GPS data on group movements and behavioral data on three of these groups (“Tenori”, “Mesas”, and “Palmas”), including regular group scans (all groups) and weekly focal follows of each individual (in Tenori group and some individuals in Mesas group). Whenever possible, groups are followed from their morning sleeping site to their evening sleeping site. The Tenori group is entirely habituated to observers on foot, and the Mesas and Palmas groups are mostly habituated. These three groups range in size from 16 (Mesas) to 17 (Tenori) to 36 individuals (Palmas). The breakdown of age / sex categories can be found in Table 1.

### Ranging data

We collected ranging data for three capuchin groups (Tenori, Mesas, and Palmas) between Jan 2018 and Apr 2019 (Fig. 3d). We spent a total of 1482 hours (220 observation days) with Tenori, 481 hours (72 days) with Mesas, and 486 hours (81 days) with Palmas. Observers recorded group locations on handheld GPS units (Garmin eTrex 10 and 20) using the “track” function, which marks a point every 10 m or 10 sec, whichever comes first. When observers lost sight of their group they turned the track function off. Location data were uploaded to Google Earth Pro version 7.3.2.5776 (Google LLC 2019) and used to create convex polygons encompassing each group’s home range. All three group range polygons contained areas not traversed by the monkeys (agricultural fields, buildings, and cattle pasture) that were excluded from the polygon area measures (Di Bitetti, 2001). The river area was conserved in ranging area, as canopy cover is generally continuous over the river and the capuchins crossed it freely.

### Vegetation survey

From Jul-Nov 2018 (wet season), we conducted a vegetation survey of the reserve to examine edge and interior forest habitats (Fig 3b-c). Edge forest was defined as forest within 100 m of a forest boundary (following (Bolt et al., 2018). Edges in this forest were created by agricultural and cattle pasture fields, land cleared for buildings, and various roads that traverse the reserve. Transects were dispersed randomly within edge (n = 20) and interior (n = 20) forest using a random number generator selecting numbers associated with points on a grid overlaid on a map of Taboga. Each 50 m transect was walked in a random direction selected by a spin of a compass bezel. If the direction selected did not allow for a full 50 m transect, then the opposite direction was chosen.

Along each transect and within 2.5 m of either side of the transect, we recorded the species and diameter at breast height (DBH) of every tree with a circumference at breast height >10 cm (FAO, 2004). We recorded canopy coverage on a scale from 1-4 (1 = 1-25% coverage; 2 = 26-50%; 3 = 51-75%; 4 = 76-100%) every meter along the transect line, first during the wet season (Jul-Nov 2018) and again during the dry season (Mar-Apr 2019).

For each transect, we calculated the following: mean DBH, mean canopy cover (wet and dry season), density (trees/m2), species richness (S, i.e., the number of tree species), and Shannon’s Diversity Index (H, which accounts for both species richness and the distribution of individuals across the species represented in the sample: Shannon & Weaver, 1949; Spellerberg, 2005). To determine whether forest edges contained more resources for capuchins and to quantify the stage of forest regeneration seen along the edges, we also categorized tree species into two groups: (1) those used by capuchins for foraging or fur rubbing, and (2) those characteristic of the early stage of successional forests (following Kalacska et al., 2004); see Table S1). Then, for each group (edge vs. interior), we calculated mean DBH, density, species richness, and Shannon’s Diversity Index.

Finally, for ease of comparison across sites, we calculated overall mean Shannon’s diversity index and Shannon’s equitability (J’, i.e., the distribution of individuals across the species in a sample: DeJong, 1975; Pielou, 1969). Shannon’s equitability ranges from 0-1, with 0 indicating an uneven distribution and 1 an equitable distribution of species (DeJong, 1975; Pielou, 1969; see Table S2).

### Vegetation survey analyses

First, we examined whether there were vegetation differences between edge and interior forest types. Because our data were not normally distributed, we used Mann-Whitney U tests to compare edge and interior transects with respect to mean DBH, density (trees/m2), species richness (S), and Shannon’s Diversity Index (H). Identical comparisons were conducted: (1) after restricting species to those used by capuchins for foraging and locomotion, and (2) after restricting species to those characteristic of the early stage of successional dry forests (Kalacska et al., 2004; see Table S1). Second, to test whether canopy cover varied between edge and interior forest and/or if canopy cover changed seasonally, we fit a linear mixed model with transect location (edge or interior), season (wet or dry), and the interaction between the two as fixed effects. We controlled for transect number as a random effect and log transformed canopy cover as the dependent variable.

### Primate survey

Between Feb-Apr 2019, we conducted a primate population survey using 32 line transects comprising pre-existing roads and paths (i.e., along canals or firebreaks) and a network of trails created by the project (19 cut trails total, each at least 0.2 km apart; Fig. 3b). Transect lengths ranged from 0.2 km to 2.2 km and we walked most transects twice (once in the morning between 6:00-10:00 and once in the afternoon between 14:00-16:00), each time in an alternate cardinal direction, for a total of 55 km in transects. Three transects were only walked once due to lack of maintenance. Transects were not surveyed when it was raining.

Transects were walked by teams of observers (typically 2 and no more than 5), traveling at a speed of 1.5 km/h and stopping every 100 m for 2 min of detailed observation (Bolt et al., 2018; Pruetz & Leasor, 2002). When more than one team searched on the same day, teams walked transects that were more than 0.2 km apart to avoid double-counting primate groups. Upon encountering a primate group (defined here as sighting one or more individuals), observers recorded the time of day, primate species, and location (using a Garmin eTrex 10 or 20 handheld GPS unit). Observers paused for 10 min to count individuals of each age/sex class, when possible, and then returned to the transect.

### Primate survey analyses

We then determined whether observations of primate groups were more likely in different forest type (i.e., edge vs. interior; Fig. 3c) or proximity to a permanent water source (i.e., <100 m vs. >100 m; Fig. 3c). For each species of monkey (i.e., capuchins, howlers), we fit a generalized linear mixed model where the dependent variable was the number of observations of primate groups for each species. We assumed the number of encounters on each transect followed a Poisson distribution whose log mean depended on forest type and proximity to water as fixed effects and transect number as a random effect. We also added a constant offset term to each model to account for different research effort on transects of different lengths.

We fit all the models with the lme4 package (Bates et al., 2015) in R version 3.3.2 (R Core Team, 2016; RStudio Team, 2016). Figures were created using the ggplot2 package (Wickham, 2009).

## RESULTS

### Capuchin density and home range size

The density of capuchins in the Taboga Forest is higher than that reported from all other long-term white-faced capuchin sites (Table 2). This high density emerges whether we use the mean group and homerange size from individual groups (21.65 individuals / km2) or the total population and total useable area (36.17 individuals / km2). For other sites, the group-based estimates range from 7.97-12.93 individuals / km2; and the total population-based estimates range from 5.60-20.00 individuals / km2 (Table 2). Therefore, with the exception of BCI, the Taboga population is 2-6 times more dense than other white-faced capuchin sites. Moreover, we believe the total estimate from the Taboga Forest is a conservative estimate because we suspect that several capuchin groups were not censused during our primate surveys (the forest continues into private land that we are not allowed to survey). We estimate that there are three additional capuchin groups here that also range into the Taboga Forest.

With respect to our habituated groups, Tenori had the smallest range size (60.5 ha, 70% of which was “edge” habitat) in comparison with Mesas (129.4 ha, 60% edge) and Palmas (102.1 ha, 57% edge) groups (Fig. 3d). We do not yet have accurate estimates of how much our capuchin groups overlap. We will soon have data from another group (Escameka group) that overlaps with the Mesas group, providing us with two points of overlap (Tenori and Palmas groups, and Mesas and Escameka groups).

### Vegetation survey

Contrary to our predictions, we found no significant differences in the interior and in the edge for mean tree DBH (Mann Whitney U; U = 185.5, *p* = 0.70; Table 3), mean tree species richness (U = 234.5, *p* = 0.37; Table 3), mean tree density (U = 203.5, *p* = 0.94; Table 3), or Shannon’s Diversity Index (U = 244, *p* = 0.24; Table 3). In comparing just trees used by capuchins, we again found no significant differences due to transect location for mean tree DBH (Mann Whitney U; U = 185, *p* = 0.70; Table 3), mean tree species richness (U = 264.5, *p* = 0.08; Table 3), mean tree density (U = 241, *p* = 0.27; Table 3), or Shannon’s Diversity Index (U = 264, *p* = 0.09; Table 3). For indicator trees, we found that trees on the edge had a significantly greater DBH than indicator trees in the interior (Mann-Whitney U; U = 274, *p* = 0.046; Table 3). We found no difference between edge and interior forest for mean indicator species richness (U = 144.5, *p* = 0.132; Table 3), mean tree density (U = 234.5, *p* = 0.36; Table 3), or Shannon’s Diversity Index (U = 153, *p* = 0.20; Table 3).

**Table 3.** Mean ± standard error of vegetation measures in the edge and interior of all trees, monkey food trees, and indicator tree species. Bold indicates significant differences between edge and interior, * denotes p<0.05.

|  | Overall |  | Monkey Food Trees |  | Indicator Trees |  |
| --- | --- | --- | --- | --- | --- | --- |
|  | *Edge* | *Interior* | *Edge* | *Interior* | *Edge* | *Interior* |
| **DBH (cm)** | 12.53 ± 1.31 | 14.30 ± 1.72 | 12.85 ± 1.31 | 15.23 ± 1.96 | **16.21 ± 1.55** | **11.30 ± 1.72 \*** |
| **Density (trees/m<sup>2</sup>)** | 0.13 ± 0.01 | 0.15 ± 0.03 | 0.08 ± 0.01 | 0.07 ± 0.04 | 0.03 ± 0.01 | 0.08 ± 0.03 |
| **Richness** | 12.30 ± 1.01 | 11.15 ± 1.23 | 7.40 ± 0.61 | 6.90 ± 1.50 | 3.00 ± 0.61 | 4.45 ± 0.72 |
| **Diversity index (H)** | 2.10 ± 0.11 | 1.89 ± 0.14 | 1.65 ± 0.12 | 1.34 ± 0.13 | 0.67 ± 0.15 | 0.99 ± 0.17 |

We found that in the dry season, interior transects had significantly less canopy cover than edge transects (Interior × Wet season; Beta = 0.42, SE = 0.17, *p* = 0.020). We found that there was more canopy coverage in the wet season months than in the dry season (GLM; Wet Season, Beta = 0.31, SE = 0.12, *p* = 0.014). We also found a significant effect of edge over the interior (Interior, Beta = −0.44, SE = 0.13, *p* = 0.001; Fig. 4). In other words, the edge forest is able to better maintain its canopy cover throughout the dry season, while the interior forest does not, following a typical deciduous pattern where trees lose their leaves during the dry season.

**Figure 4.**
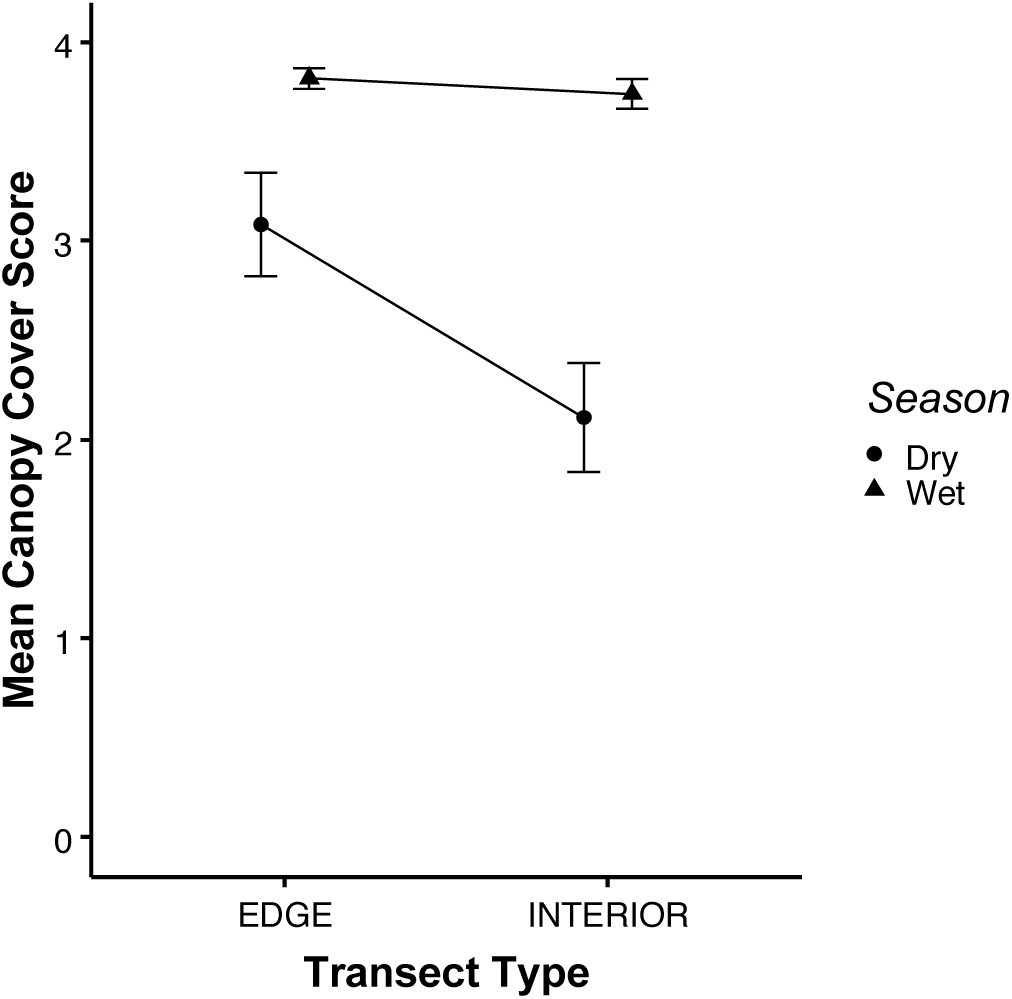
Mean canopy cover score (± standard error) across season and transect type.

### Population survey

As predicted, both capuchins and howlers showed neutral edge effects (i.e., no significant difference between group encounter rates in edge vs. interior forest). Capuchin encounter rates were lower overall (compared to howler encounters) and did not differ between edge (0.34 groups/km; CI: −2.10, 0.48) and interior forest (0.46 groups/km; CI: −0.48, 2.10; *p* = 0.25; Fig. 5a). Although there was a higher encounter rate for howlers in edge (1.31 groups/km; CI: −0.38, 0.85) compared to interior forest (0.75 groups/km; CI: −0.85, 0.38; Fig. 5c), this difference was not significant (*p* = 0.43).

**Figure 5a-d.**
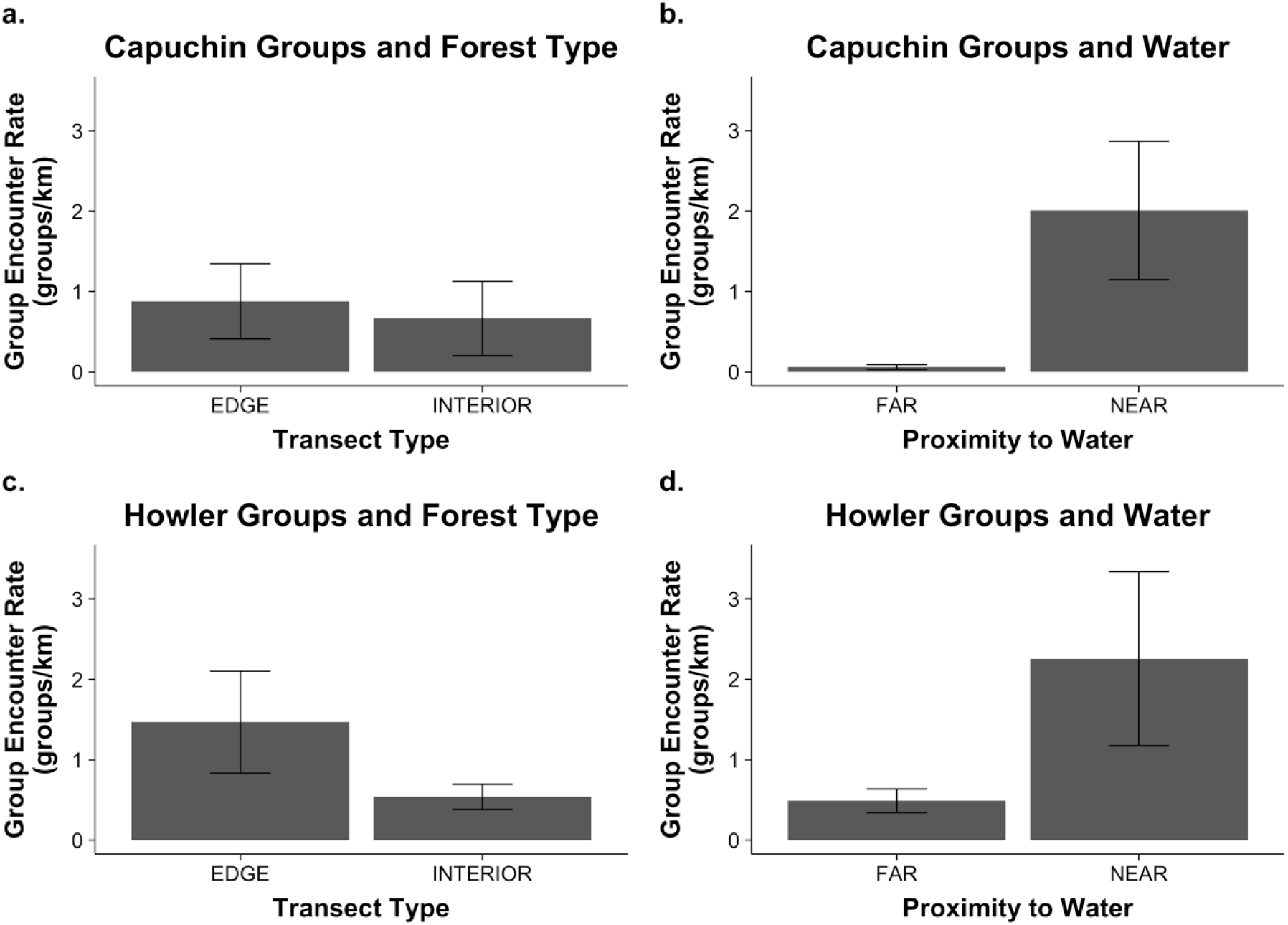
Mean group encounter rate (groups/km walked ± standard error) by forest type in Taboga. **(a)** capuchin groups in edge and interior forest; **(b)** capuchin groups near (<0.01 km) or far (>0.01 km) from a permanent water source; **(c)** howler groups in edge and interior forest; **(d)** howler groups near and far from a permanent water source.

As predicted, capuchin group encounters were higher near permanent water sources (i.e., the river or large canals: 0.78 groups/km; CI: 1.42, 4.41) compared to farther from water (0.15 groups/km; CI: −4.41, −1.42; *p* = 1.73 × 10-4; Fig. 5b). Contrary to our predictions, however, howler group encounter rates were also significantly higher near water sources (1.69 groups/km; CI: 0.42, 1.72) compared to farther from water (0.57 groups/km; CI: −1.72, −0.42; *p* = 1.24 × 10-3; Fig. 5d).

## DISCUSSION

The Taboga Forest of Costa Rica has one of the highest densities of white-faced capuchins thus far recorded. Here, we asked whether specific features of the forest might allow these capuchins to survive and reproduce at such high densities. Our results suggest that the presence of reliable year-round water sources is critical for capuchins (and possibly for howlers) living in a seasonally dry habitat. For example, capuchins in Santa Rosa National Park (another tropical dry forest in Costa Rica) rely on a limited number of water holes during the dry season, and access to these water holes is thought to be the primary constraint on the capuchin population (Fedigan & Jack, 2001; Fedigan, Rose, & Avila, 1996, 1998). In contrast, Taboga has two types of year-round water supply: the river and a system of canals. We did not test whether forest characteristics varied significantly according to distance from water sources. However, because the canals have cement bottoms, we think that it is unlikely that the canals impact the surrounding flora all that much. We will implement future studies to test how forest characteristics vary with proximity to the river (and the associated riparian/semi-deciduous forest type). For example, riparian forests may contain certain fruiting trees central to the capuchin diet, overall larger trees due to year-round water supply, and/or year-round canopy cover. Our results also suggest that howlers at Taboga may be more dependent on permanent water sources than at other sites, as they were also frequently found near permanent water sources. However, this may have more to do with the forest subtype near the river (i.e., evergreen and riparian) than the need to drink water daily (Fedigan & Jack, 2001).

In line with previous research that found neutral edge effects for both species (e.g., (Bolt et al., 2018), we found no difference between capuchin (or howler) group encounter rates when we compared edge vs. interior forest. Combined with the overall high capuchin population density, this suggests that despite a large percentage of edge forest (nearly 40% of the 516 hectares), capuchins appear to thrive in forest fragments (Cunha et al., 2006). Indeed, we found that capuchins were equally likely to find staple food and fur-rubbing species in the edge compared to the interior forest and that the size of these staple species (i.e., DBH) did not vary significantly between edge and interior. Other features of the forest, like canopy height (Fleagle & Mittermeier, 1980) and canopy cover (Fedigan & Jack, 2001) have been useful in explaining forest use by other primate taxa. Although we did not record canopy height in this study, we found that the DBH for our trees did not differ from the edge to the interior. Canopy cover showed a very different pattern though. The edge forest in Taboga maintained canopy cover even throughout the dry season, while the interior forest was more deciduous (we expand on possible reasons for this below). For primates, semi-evergreen forest can provide shade and may stay cooler through the hottest months (Fedigan & Jack, 2001; Fedigan et al., 1996), and therefore both capuchins and howlers might spend more time in edge forest during the dry season (when our primate survey took place) than they do during the wet season. Longitudinal data will determine whether ranging patterns vary seasonally.

Together, our data suggest that the difference between edge and interior forest at Taboga is less than that from other sites (e.g., Arroyo-Rodríguez & Mandujano, 2009; Bolt et al., 2018; Harris, 1988; Lehman, Rajaonson, & Day, 2006; Saunders, Hobbs, & Margules, 1991). This may be because the initial anthropogenic disturbance (i.e., creation of pastures and croplands around the reserve) happened some time ago and the forest is actually in the intermediate stages of regeneration (Kalacska et al., 2004). Three lines of evidence support this hypothesis. First, both edge and interior forest at Taboga exhibit high species richness and diversity, which also characterize intermediate tropical dry forest succession at nearby Santa Rosa (Kalacska et al., 2004). Second, tree species that characterize the first stage of tropical dry forest succession (of which many remain through stages 2 & 3) are well-established in the forest edge. Specifically, these trees had a significantly higher mean DBH in edge forest compared to interior forest. Third, the early stages of dry forest succession are characterized by a high percentage of deciduous trees (Kalacska et al., 2004). Yet, we found that the edge forest in Taboga was semi-evergreen throughout the dry season. This, of course, raises the question of why the interior forest is more deciduous. It may be that the Taboga Forest is small enough that it lacks a true “interior” (Banks-Leite, Ewers, & Metzger, 2010), and therefore the entire forest represents different stages of regeneration. Alternatively, much of the interior forest is also more elevated and may lack year-round water sources. Finally, flood-irrigation of agricultural land during the dry season might spill-over into edge forests, thus allowing for year-round canopy maintenance.

Our survey indicates that the Taboga Forest is actually composed of at least three distinct forest types: (1) the deciduous tropical dry forest that extends from the North boundary of the farm down to the river, (2) the riparian semi-deciduous forest that follows the river, and (3) a moist palm forest that retains canopy cover year-round. Future studies should distinguish between these subtypes to test how different microclimates alter edge effects. In addition, the severity of edge effects on forest composition can vary according to the type of disturbance (i.e., road, pastureland, sugar cane/rice plantation, etc.). Thus, the universal 100 m buffer used here may not apply equally to each forest type found in Taboga (Didham et al., 2015; Harper et al., 2005). For capuchins living in a highly seasonal environment, having distinct habitats may provide a buffer from extreme fluctuations in temperature and rainfall (indeed, preliminary data suggest group ranging varies significantly by forest type between seasons). For example, in 2010, a capuchin population crash on Barro Colorado Island was caused by unusually heavy rains that decimated the arthropod population (a key source of protein for capuchins) (Milton & Giacalone, 2014). In Taboga, the wet and evergreen forest may be less vulnerable to drought or El Niño events (e.g., Campos, Jack, & Fedigan, 2015), while the deciduous tropical dry forest may be less vulnerable to unusually heavy rains and flooding.

The abundance of the capuchins in Taboga has important implications for conservation efforts. For certain species, the size and disturbance of a forest fragment may matter less than the composition and availability of key resources, like above-ground water. Our analysis here adds to our understanding of factors that influence primate abundance, and also establishes Taboga as critical case study in tropical dry forest dynamics. Future studies will provide a more fine-grained analysis of the possible interaction between edge effects, habitat type, and season, and how these factors influence primate sightings (Gogarten et al., 2012). For example, we were not able to test here whether primates prefer the river over human-made canals (or vice-versa), though we predict that howlers sightings may be more frequent along the river (i.e., that howlers prefer riverine habitat over others but do not necessarily need to be close to an above-ground water source). For capuchins, the next question is how the high density in Taboga influences ranging patterns, home range overlap, and the frequency and intensity of intergroup encounters (Perry, 1996; Perry 2012). Preliminary data suggest that intergroup encounters are higher at Taboga than at other sites, but that the intensity of such encounters is lower, which may represent a behavioral adaptation to frequent encounters.

## ACKNOWLEDGEMENTS

We are grateful to the following Costa Rican institutions for allowing us to work in the Taboga Forest since 2017: the Ministerio de Ambiente y Energía (MINAE), the Comisión Nacional para la Gestión de la Biodiversidad (CONAGEBIO), and the Universidad Técnica Nacional (UTN). We would also like to thank the following people: Dr. Susan Perry for guidance and support; Courtney Anderson and Jahmaira Archbold for skilled field assistance during our first year; and the students of UTN who assisted with data collection. Finally, we would like acknowledge the University of Michigan for their financial support. The authors have no conflicts of interest to declare.

