## Supplementary material for "High density of white-faced capuchins (*Cebus capucinus*) and habitat quality in the Taboga Forest of Costa Rica": Table S1

**Table S1.** List of tree species identified overall, included as relevant to capuchins specifically, and included as representative of the early stage of successional tropical dry forest regeneration.

| | **Vegetation Survey Species** | |
|---|---|---|
| *All Species (Non-native species in bold)* | *Species utilized by Cebus capucinus for alimentation or fur rubbing* | *Species characteristic of early stage successional tropical dry forest* |
| *Acacia collinsii* | X | X |
| *Acacia cornigera* | X | |
| *Acacia farnesiana* | | |
| *Acacia tenuifolia* | | |
| *Acosmium panamense* | | |
| *Albizia adinocephala* | | |
| *Albizia niopoides* | | |
| *Albizia saman* | X | |
| *Alibertia edulis* | | |
| *Alibizia guachapele (Pseudosamanea guachapele)* | X | |
| *Allophylus racemosus* | X | |
| *Alvaradoa amorphoides* | | |
| *Anacardium excelsum* | X | |
| *Annona purpurea* | X | |
| *Annona reticulata* | X | |
| *Apeiba tibourbou* | | X |
| *Ardisia revoluta* | X | X |
| *Attalea rostrata* | X | |
| *Bactris guianeensis* | X | |
| *Bauhinia pauletia* | | |
| *Bixa urucurana* | X | |
| *Brosimum alicastrum* | X | |
| *Bursera simaruba* | X | X |
| *Byrsonima crassifolia* | X | X |
| *Calycophyllum candidissimum* | | |
| *Capparis baducca* | | |
| *Capparis indica* | | |

| | | |
|---|---|---|
| *Casearia aculeata* | X | |
| *Casearia arguta* | X | X |
| *Casearia corymbosa* | | |
| *Casearia sylvestris* | X | |
| *Cecropia peltata* | X | X |
| *Cedrela odorata* | | X |
| *Chomelia spinosa* | X | |
| ***Citrus limon*** | X | |
| *Coccoloba caracasana* | X | |
| *Cochlospermum vitifolium* | | X |
| *Cordia alliodora* | | X |
| *Cordia collococca* | | |
| *Cordia dentata* | X | X |
| *Cordia panamensis* | X | |
| *Crescentia cujete* | | |
| *Critonia moriflora* | | |
| *Dalbergia retusa* | | X |
| *Diospyros salicifolia* | X | X |
| *Enterolobium cyclocarpum* | | X |
| *Erythroxylum havanense* | X | |
| *Eugenia oerstediana* | X | |
| *Eugenia salamensis* | X | |
| *Exostema mexicanum* | | |
| *Genipa americana* | X | X |
| ***Gmelina arborea*** | X | |
| *Godmania aesculifolia* | | |
| *Guarea glabra* | | |
| *Guazuma ulmifolia* | X | X |
| *Gyrocarpus jatrophifolius* | | |
| *Inga vera* | X | |
| ***Leucaena leucocephala*** | | |
| *Licania arborea* | | |
| *Lonchocarpus felipei* | | |

| | | |
|---|---|---|
| *Lonchocarpus guatemalensis* | | |
| *Lonchocarpus minimiflorus* | | |
| *Lonchocarpus parviflorus* | | |
| *Luehea candida* | X | X |
| *Luehea seemannii* | | |
| *Luehea speciosa* | | X |
| *Lysiloma divaricatum* | | |
| *Machaerium biovulatum* | | X |
| *Maclura tinctoria* | X | |
| *Malvaviscus arboreus* | X | |
| ***Mangifera indica*** | X | |
| *Manilkara chicle* | | |
| *Margaritaria nobilis* | X | |
| *Myrospermum frutescens* | | |
| *Nectandra membranacea* | | |
| *Ocotea veraguensis* | | |
| *Pachira quinata* | X | |
| *Picramnia antidesma* | | |
| *Piper reticulatum* | X | |
| *Piper tuberculatum* | X | |
| *Piscidia carthagenensis* | | |
| *Pisonia aculeata* | | |
| *Pithecellobium lanceolatum* | | X |
| *Plumeria rubra* | | |
| *Pricamnia antidesma* | | |
| *Psychotria carthagenensis* | X | |
| *Ruprechtia costaricensis* | | |
| *Sapindus saponaria* | | |
| *Schizolobium parahyba* | | |
| *Semialarium mexicanum* | | X |
| *Senna papilosa* | | |
| ***Senna siamea*** | | |
| *Sloania terniflora* | | |

| | | |
|---|---|---|
| *Solanum hazenii* | X | |
| *Spondias mombin* | X | X |
| *Spondias purpurea* | X | X |
| *Spondias radlkoferi* | | |
| *Stemmadenia obovata* | X | |
| *Sterculia apetala* | X | |
| *Swietenia macrophylla* | | X |
| *Tabebuia ochracea* | X | X |
| *Tabebuia rosea* | | |
| *Tabernaemontana alba* | | |
| *Tabernaemontana donnell-smithii* | X | |
| ***Tectona grandis*** | | |
| *Terminalia oblonga* | | |
| *Trichilia americana* | | |
| *Trichilia martiana* | X | |
| *Trichilia trifolia* | | |
| *Triplaris melaenodendron* | | |
| *Trophis racemosa* | | |
