## Supplementary material for "High density of white-faced capuchins (*Cebus capucinus*) and habitat quality in the Taboga Forest of Costa Rica": Table S2

**Table S2.** Comparison of forest characteristics across related sites.

| Site | Mean Density ($m^2$) | Species Richness (S) | Shanon's Diversity Index (H) | Shanon's Equitability (J') |
|---|---|---|---|---|
| **Tropical Dry Forest, Costa Rica** | | | | |
| Taboga: *Overall* | | 111 | 3.84 | 0.82 |
| Taboga: *Edge* | 0.13 | 12.30 | 2.10 | |
| Taboga: *Interior* | 0.15 | 11.5 | 1.89 | |
| Palo Verde National Park (Gillespie et al. 2000, Powers, et al. 2009) | 0.227 | 65 | 1.75 | |
| Santa Rosa National Park: Overall (Gillespie et al., 2000) | 0.246 | 75 | | |
| Santa Rosa National Park: *Early successional stage* (Kalacska et al. 2007) | | | 1.77 | |
| Santa Rosa National Park: *Intermediate successional stage* (Kalacska et al. 2007) | | | 2.88 | |
| Santa Rosa National Park: *Late successional stage* (Kalacska et al. 2007) | | | 2.75 | |
| **Tropical Lowland Rainforest, Costa Rica** | | | | |
| La Suerte Biological Research Station: *Edge* (Bolt et al. 2018) | 0.11 | 3.8 | | |
| La Suerte Biological Research Station: *Interior* (Bolt et al. 2018) | 0.08 | 6.1 | | |
| **Tropical/Subtropical Dry Forest, Mexico** | | | | |
| Nizanda: *40 years after initial abandonment* (Lebrija-Trejos et al. 2008) | 0.7475[+] | 61 | 3.34 | 0.71-0.77 |
| Nizanda: *Mature forest* (Lebrija-Trejos et al. 2008) | 0.246 | 112 | 2.9 | 0.91 |

[+]Originally reported as individuals/ha

Full citations:
Bolt, L. M., Schreier, A. L., Voss, K. A., Sheehan, E. A., Barrickman, N. L., Pryor, N. P. & Barton, M. C. (2018). The influence of anthropogenic edge effects on primate populations and their habitat in a fragmented rainforest in Costa Rica. *Primates*, *59*:301-311.

Gillespie, T. W., A. Grijalva, and C. N. Farris (2000). Diversity, composition, and structure of tropical dry forests in Central America. *Plant Ecology, 147*:37-47.

Kalacska, M., G. A. Sanchez-Azofeifa, B. Rivard, T. Caelli, H. P. White, and J. C. Calvo-Alvarado. 2007. Ecological fingerprinting of ecosystem succession: estimating secondary tropical dry forest structure and diversity using imaging spectroscopy. Remote Sensing of Environment 108: 82-96. doi: 10.1016/j.rse.2006.11.007

Lebrija-Trejos, E., F. Bongers, E. A. Pérez-García, and J. A. Meave. (2008). Successional change and resilience of a very dry tropical deciduous forest following shifting agriculture. *Biotropica, 40*:422-431.

Powers, J. S., J. M. Becknell, J. Irving, and D. Pérez-Aviles. 2009. Diversity and structure of regenerating tropical dry forests in Costa Rica: geographic patterns and environmental drivers. Forest Ecology and Management 258: 959-970. doi: 10.1016/j.foreco.2008.10.036
